# One-week escitalopram intake shifts excitation-inhibition balance in the healthy female brain

**DOI:** 10.1101/2021.07.09.451806

**Authors:** Rachel G. Zsido, Eóin N. Molloy, Elena Cesnaite, Gergana Zheleva, Nathalie Beinhölzl, Ulrike Scharrer, Fabian A. Piecha, Ralf Regenthal, Arno Villringer, Vadim V. Nikulin, Julia Sacher

## Abstract

Neural health relies on cortical excitation-inhibition balance (EIB), with disrupted EIB underlying circuit dysfunction in several neuropsychiatric disorders. Previous research suggests links between increased cortical excitation and neuroplasticity induced by selective serotonin reuptake inhibitors (SSRIs). Whether there are modulations of EIB following SSRI-administration in the healthy human brain, however, remains unclear. To this end, we assessed changes in EIB following longitudinal escitalopram-intake. In a randomized, double-blind study protocol, a sample of 59 healthy female individuals on oral contraceptives underwent three resting-state electroencephalography recordings after daily administration of 20 mg escitalopram (n = 28) or placebo (n = 31) at baseline, after single dose, and after 1 week (steady state). We assessed 1/f slope of the power spectrum, a marker of EIB, compared individual trajectories of 1/f slope changes contrasting single dose and 1-week drug intake, and tested the relationship of escitalopram plasma levels and cortical excitatory and inhibitory balance shifts. Escitalopram-intake associated with decreased 1/f slope, indicating an EIB shift in favor of excitation. Furthermore, 1/f slope at baseline and after single dose of escitalopram predicted 1/f slope at steady state. Higher plasma escitalopram levels at single dose associated with better maintenance of these EIB changes throughout the drug administration week. Characterizing changes in 1/f slope during longitudinal SSRI-intake in healthy female individuals, we show that escitalopram shifted EIB in favor of excitation. These findings demonstrate the potential for 1/f slope to predict individual cortical responsivity to SSRIs and widen the neuroimaging lens by testing an interventional psychopharmacological design in a clearly-defined endocrinological state.

## 1. Introduction

The balance of excitation and inhibition in neuronal circuits is essential for brain network function and stability (1, 2). Evidence shows that failure to maintain this excitation-inhibition balance (EIB) can underlie circuit dysfunction observed in several neuropsychiatric and neurodevelopmental disorders (3), such as autism (4, 5), schizophrenia (5–7), and major depressive disorder (8). Conceptual models propose that selective serotonin reuptake inhibitors (SSRIs), a class of antidepressants upregulating serotonergic transmission, act by enhancing synaptic plasticity (9–11). Findings from animal studies (12, 13) suggest that alterations in cortical excitation and inhibition may be a critical factor driving SSRI-induced plasticity. While many of these findings are limited to animal models, some studies have investigated SSRI-induced changes in EIB in human participants (14–16). However, most of these studies investigated only a single kinetic state (e.g., a single dose of the drug) or relatively small samples. Given the widespread use and highly variable response rates to SSRIs (17, 18), it is of clear clinical interest to understand both the influence of longitudinal SSRI-intake on EIB and to identify a neurophysiological marker of EIB that could predict individual cortical responsivity to SSRIs.

Resting-state electroencephalography (rs-EEG) provides a reliable, non-invasive method for investigating pharmacologically-driven alterations in human cortical activity. Power of alpha oscillations could provide insight into EIB alterations due to its functional role in cortical inhibition (19). For example, decreases in relative alpha power, thought to reflect enhanced cortical excitability (19, 20), have been observed in healthy male participants (n = 12) following one week of litoxetine administration, an SSRI under development (21), as well as in depressed patients following one week of escitalopram administration (22). Another exploratory study in healthy male participants (23) (n = 14/group) found that decreased serotonin synthesis via tryptophan depletion was associated with a trend towards increased relative alpha power. A systematic review of the effects of SSRIs in healthy participants reported decreases in power of alpha oscillations following a single administration of low and medium dose SSRIs (24); alpha power results for high doses, however, were inconclusive (24). Given that (1) alpha power is associated with inhibitory processes, (2) serotonergic manipulation has been shown to modulate alpha power, and (3) abnormal alpha power is associated with psychiatric symptoms in clinical populations such as depression (25, 26), it is possible that SSRIs may act via decreases in alpha-band activity that reflect a shift in favor of cortical excitation in the human brain.

Beyond the more canonically-defined measures such as alpha power, 1/f slope of the power spectral density (PSD) is thought to more directly reflect EIB. Neurophysiological brain signal consists of periodic oscillatory activity and aperiodic activity (1/f slope) of the non-oscillatory PSD background (27), which have been shown to play functionally distinct roles (28). While more conventional approaches have focused on narrowband oscillations or frequency band ratios, there has been a current upsurge in neuroscience research in the past year focusing on 1/f slope as a unique neurophysiological marker (6, 28–31). Simulation data with local field potentials demonstrate that 1/f slope inversely reflects EIB (32), a finding that has been validated in *in vivo* animal models in which anesthesia administration led to an increase in 1/f slope or steepening of the PSD decay (32). A recent interventional rs-EEG study in healthy humans showed that ketamine and anesthesia administration, which respectively tip the balance in favor of cortical excitation (increase in EIB) and inhibition (decrease in EIB), resulted in the expected decrease and increase in 1/f slope (33). Since then, several recent studies have shown the reliability of this measure using human scalp EEG activity in both healthy (29–31) and clinical populations (6, 29, 34).

Nevertheless, the effect of SSRIs on 1/f slope has yet to be investigated. To clarify the neurophysiological mechanisms underlying serotonergic action in health and identify a potential marker to predict individual cortical responsivity to SSRIs, we require a longitudinal model of how SSRI-intake affects EIB in a homogenous sample of healthy controls. In this study, we administered a clinically-relevant dose of 20 mg escitalopram for seven days (35–39). Given known sex differences in neural responses to serotonergic intervention (40, 41), higher antidepressant prescription rates in women (42) as well as the urgent need to increase female samples in neuroscience research (43–45), we recruited 59 healthy female participants (28 escitalopram, 31 placebo) who underwent rs-EEG at 3 time-points: before randomization (baseline), after a single dose of administration (single dose), and after one week of daily administration (steady state) (35). All participants were using oral contraceptives to downregulate ovarian hormone fluctuations to control for potential effects of sex hormones on escitalopram responsivity (40, 46), brain resting-state connectivity (45, 47–49), and resting-state alpha activity specifically (50). We estimated 1/f slope of the PSD as a measure of EIB. Given our decision to separately investigate oscillatory activity from 1/f activity, we calculated power of alpha oscillations independently from aperiodic activity. To compare to previous findings, however, we additionally calculated relative alpha power from the original PSD. We hypothesized that escitalopram administration would be associated with decreases in both 1/f slope and power of alpha oscillations.

## 2. Materials and Methods

### 2.1 Participants and eligibility

Participants provided written informed consent after study procedures were explained. Eligible individuals were female, right-handed, 18-35 years old, with a body mass index (BMI) between 18.5-25 kg/m^2^, and without any neurological or psychiatric illness as confirmed with a structured clinical interview (51, 52). All participants were taking oral contraceptives for ≥ 3 months to downregulate ovarian hormone fluctuations (53). Exclusion criteria were medication, tobacco or alcohol use, positive drug or pregnancy tests, and abnormal QT times in electrocardiogram readings. Of the 87 participants screened, 70 were enrolled. We included 59 participants in analyses as 6 (4 escitalopram) chose to discontinue and 5 (3 escitalopram) were excluded after data quality assessment. Participants were under medical supervision for the entire experiment. The Ethics Committee of the Faculty of Medicine of Leipzig University (approval # 390/16-ek) approved all procedures.

### 2.2 Study design and experimental protocol

These data were acquired as a secondary outcome measure from a randomized, double-blind, parallel study design (ClinicalTrials.gov: NCT03162185, Open Science Framework https://osf.io/g9usb), as previously reported (54). The present study was designed to test the hypothesis that one-week SSRI-administration shifts cortical EIB using a novel EEG surrogate marker (32), and to investigate whether the acute EIB response to a single dose of escitalopram can predict individual EIB responsivity after 7 days of drug-intake (when plasma levels no longer fluctuate and relative steady state levels in healthy participants are reached (35). We administered 20 mg of escitalopram, which reliably blocks 80 percent of serotonin transporter and achieves steady state conditions after one week of administration (35–39)) or an identical placebo capsule (mannitol/aerosol) from sequentially numbered containers at fixed times each day for seven consecutive days. We recorded a baseline rs-EEG prior to drug administration (baseline). Participants were then randomized to receive either escitalopram or placebo. Randomization employed an independent block randomization with a 1:1 allocation ratio, conducted by the Pharmacy of the University Clinic at Leipzig University. The experimenter and participants were blind to treatment allocation. We recorded another rs-EEG measurement following a single dose (single dose) and after seven days of administration (when relatively steady state plasma levels are reached) (**Figure 1**).

**Figure 1.**
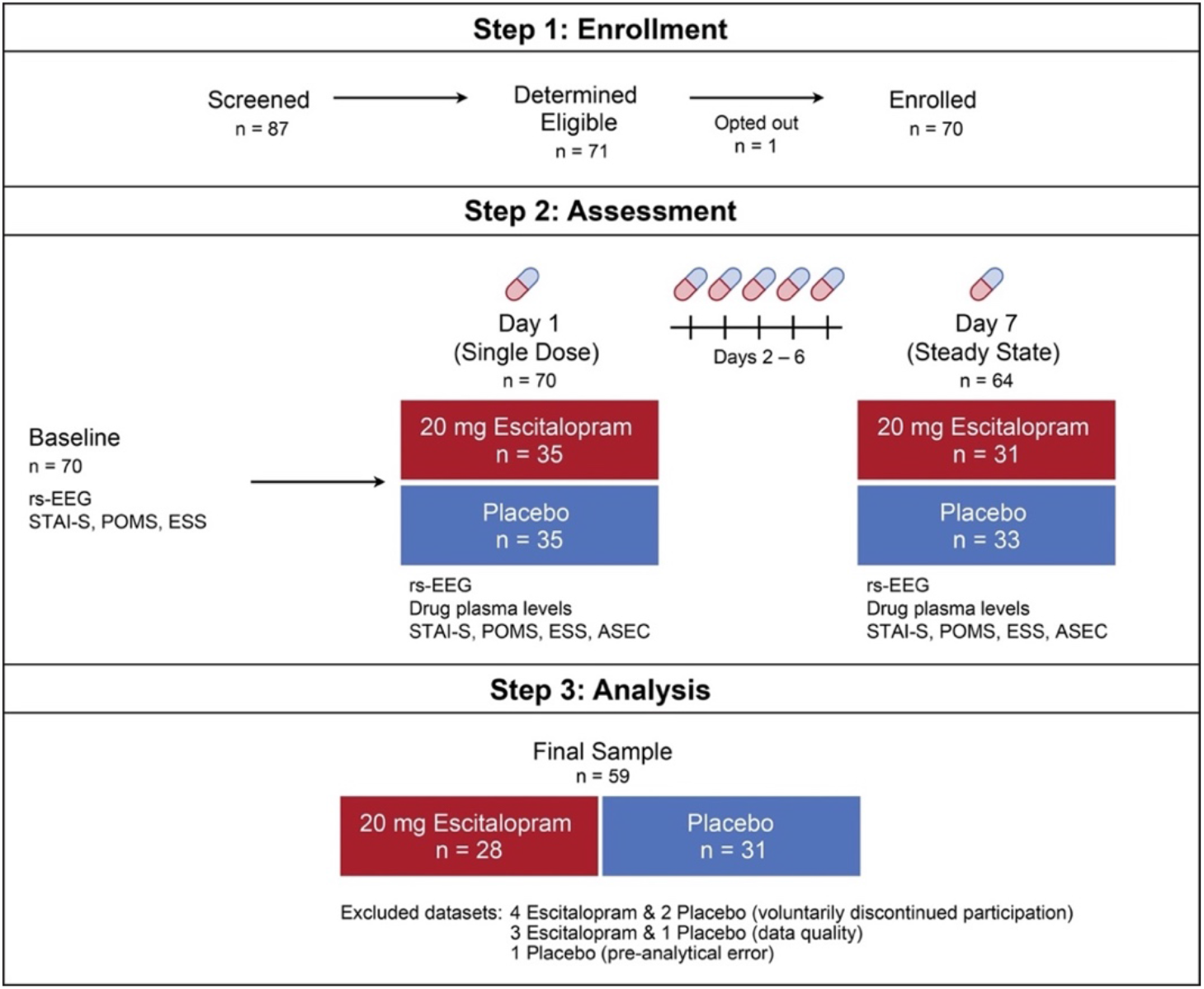
Study design and experimental protocol. Step 1 details screening and enrollment numbers. Step 2 depicts study design, with either escitalopram or placebo administered for seven days following a baseline resting-state electroencephalography (rsEEG) recording. In total, 6 participants voluntarily discontinued participation during this phase. Step 3 lists the final sample included in the analyses (n = 59). State-Trait Anxiety Inventory (STAI-S), Profile of Mood States (POMS), Epworth Sleepiness Scale (ESS), Analysis of Antidepressant Side-Effect Checklist (ASEC).

All rs-EEG measurements took place at approximately the same time of day for participants and 4-5 hours after escitalopram or placebo intake, as previous pharmacokinetic modeling in healthy participants suggests maximum drug plasma concentration is reached 3-5 hours after 20 mg oral escitalopram intake (55, 56). We assessed potential changes in anxiety (State-Trait Anxiety Inventory, STAI-S (57, 58), non-patient edition), current mood (German version of Profile of Mood States, POMS (59)), daytime sleepiness (Epworth Sleepiness Scale, ESS (60)) and escitalopram side effects (Antidepressant Side-Effect Checklist, ASEC (61)). The STAI, POMS, ESS, and ASEC took place at approximately the same time of day for each participant. Serum follicle-stimulating hormone (FSH) and luteinizing hormone (LH) levels were measured by electrochemiluminescence immunoassay (ECLIA) (Cobas; Roche) prior to enrollment to confirm downregulated ovarian hormones. We analyzed plasma escitalopram concentration from single dose and one-week steady state using a validated high-performance liquid chromatography method (62), with four quality control samples covering the low, medium, and high range of the calibration curve. Deviation of the measured concentrations of the quality control samples was tested for an acceptance interval of ± 15%.

### 2.3 rs-EEG acquisition

We used a 32-channel EASYCAP (Brain Products GmbH, Germany) electrode cap, BrainAmp amplifier, and Brain Vision Recorder (Brain Products GmbH, Germany). Sintered Ag/AgCl point electrodes were mounted using the international 10-20 system (63), and impedance levels per electrode were maintained at < 10 kΩ (typically < 5 kΩ). The reference (M2) electrode was placed on the right mastoid and an additional electrode (M1) was placed on the left mastoid. Four electrodes were placed to monitor eye movement and one ground electrode was placed on the sternum. Data were recorded using a sampling rate of 1 kHz, a high-pass filter of 0.015 Hz, and a low-pass filter of 250 Hz. Participants sat in an acoustically-shielded room with eyes closed for 11 minutes, with a 30-second break after 5.5 minutes.

### 2.4 rs-EEG preprocessing

Data were preprocessed using EEGLAB toolbox (v14.1) in MATLAB (v9.3). EEG data were band-pass filtered between 1-45 Hz (4th order, forward and backward directions, Butterworth filter) and a notch filter was applied at 50 Hz to ensure artifact removal related to power line noise. Data were down-sampled to 500 Hz. The 30-second break was removed, creating a single 11-minute recording. Bad segments from the time series data were marked and rejected by an algorithm with individually adjusted noise thresholds: for low frequencies (1-15 Hz), the threshold was set to 3 standard deviations above the filtered mean amplitude; for higher frequencies (15-45 Hz), the threshold was 40 μV. Electrocardiogram, electrooculography, and two frontal channels (Fp1, Fp2) were removed prior to bad segment estimation due to high amplitudes of eye blinks and heartbeat artifacts. Marked bad segments were applied to the full dataset of 27 scalp electrodes. Flagged bad segments (> 60 seconds) were manually reviewed while broken channels were assessed with visual inspection of the PSD and excluded if necessary. Bad segments were removed prior to re-referencing to ensure noise was not projected to other channels, and so that independent component analysis (ICA) is performed on data that is not contaminated by major noise artifacts spread over all electrodes (64). We re-referenced to the common average, where every electrode is referenced against the average of all electrode recording, to avoid prioritizing voltage differences coming from one specific location. ICA weights (Infomax (65)) were then calculated on remaining segments of the time series for each participant that were used to project out ocular, muscular, and cardiac activity components. The second mastoid electrode (M2) was then removed.

### 2.5 Rs-EEG data analysis

We estimated 1/f offset and slope per channel from the PSD (Welch’s PSD with 4-second windows overlapping by 50%) of the preprocessed signal in a frequency range of 1-40 Hz using the FOOOF toolbox (27) in Python (v3.5), a module for parameterizing neural power spectra that quantifies both the periodic and aperiodic activity from the PSD (27). Major oscillatory peaks were excluded when estimating slope of 1/f decay. This allowed detrending of the PSD by subtracting the estimated non-logged 1/f decay from the original PSD. Individual alpha peak frequency was measured on a detrended PSD by a peak-search in a range between 7-13 Hz. A peak was defined as a curve exceeding a threshold of 0.05 μV2/Hz. If several peaks were found, we considered the largest one. Taking individual alpha peak frequency as an anchor point at which the peak maxima appeared, we defined the range (no more than +/- 3 Hz from maxima of the peak) and used it to estimate alpha power that was defined as a summed area under the detrended PSD curve. No alpha peak was found in 4 participants (3 escitalopram) in ≥ 10 channels, thus they were excluded from analyses. Due to non-normal distribution, alpha power values were log-transformed. To detect possible outliers, we computed a mean over channels per participant at each assessment for 1/f slope and alpha power, using a cutoff of +/- 3 standard deviations within each group. No outliers were detected.

### 2.6 Statistical analysis

#### 2.6.1 Monitoring

We assessed potential group differences in age, BMI, and endogenous hormonal profiles at baseline using independent samples *t*-tests in R (v3.5.2) (66). We assessed potential group differences in total ASEC, POMS, STAI-S, and ESS scores at both single dose and one-week steady state. Questionnaire results were considered statistically significant at a Bonferroni-corrected threshold of *p* < 0.006 to account for multiple comparisons. For questionnaires that showed significant group differences, we conducted bivariate correlational analyses to test for potential associations between total questionnaire scores and either escitalopram plasma levels, alpha power, or 1/f slope.

#### 2.6.2 Linear mixed effects modeling

We analyzed mean rs-EEG 1/f slope and alpha power using a random-intercept mixed effects modeling approach. We applied one model to each metric using the ‘lmer’ function in the ‘lme4’ R package, with group and time as factors and specific outcome as dependent variable. We compared the contribution of each fixed main effect and interaction term in an omnibus modelling approach using a chi-square log-likelihood ratio test. Model contributions for each level of the fixed effects were determined using marginal *R^2^* change. Post-hoc Wilcoxon rank-sum tests and Wilcoxon signed-rank tests, implemented with the ‘wilcox.test’ function, were conducted on mean whole-brain signal.

#### 2.6.3 Cluster-based permutation tests

For significant outcomes derived from linear mixed effects modeling, we performed cluster-based permutation tests to assess spatially-specific effects of escitalopram. Given the non-normal distributions of parameters derived from EEG data, we applied non-parametric Wilcoxon rank-sum tests and Wilcoxon signed-rank tests in MATLAB to test between and within-group effects. *Z*-values obtained per electrode were then used in cluster statistics (67). Significant electrode clusters were defined as ≥ 2 neighboring electrodes significant at *p* < 0.05. The most robust cluster was validated with permutation tests (n = 1000). Briefly, the original cluster derived from the data was compared to the clusters formed by randomized partitions (i.e., randomized group information for between-group, day information for within-group) and running the same statistical tests. Next, we calculated the proportion of random significant clusters (n = 1000) that result in a larger statistic than the originally observed one; this is the cluster-level p-value, and the observed cluster is significant if *p* < 0.05.

#### 2.6.4 Linear regression on 1/f slope

We tested if 1/f slope at one time predicted 1/f slope at a later time in the escitalopram group. We conducted three regressions in R (baseline to single dose, baseline to one-week steady state, single dose to one-week steady state) using whole brain unstandardized residuals to control for trait 1/f slope signal.

#### 2.6.5 Moderation analysis

In the escitalopram group, we tested regression pathways in an exploratory moderation analysis using PROCESS macro (v3.5.3 SPSS) (68), a program using an ordinary least squares-based path analytical framework to test direct/indirect associations. We assessed significance and stability of the interaction of single dose plasma escitalopram levels (moderator) and residual single dose 1/f slope (controlled for baseline, independent variable) in association with residual one-week steady state 1/f slope (outcome variable). Variables were mean-centered, and we implemented a 95% bias-corrected bootstrap CI (BBCI), excluding 0 and based on 10,000 bootstrap samples to account for a non-normal data distribution in 1/f slope.

#### 2.6.6 Conventional analysis with relative alpha power

We calculated relative alpha power on the non-detrended (‘conventional’) PSD to compare approaches. Relative alpha power was calculated by dividing alpha-band power of individually pre-defined alpha range (procedure described above) by the total spectral power (3-45 Hz).

## 3. Results

### 3.1 Monitoring

Analyses included 59 participants (28 escitalopram, 31 placebo). We did not observe group differences for any demographic characteristics (**Table 1**). LH and FSH values were within reference range expected for downregulated hormone profiles (ECLIA, Cobas:Roche). Plasma drug levels of all participants in the escitalopram group at day 7 achieved expected steady state plasma levels (Mean±Standard Deviation 45.33±11.26 ng/mL, range 26.6-66.3) based on previously reported therapeutic reference ranges of steady-state plasma concentrations.

**Table 1.**
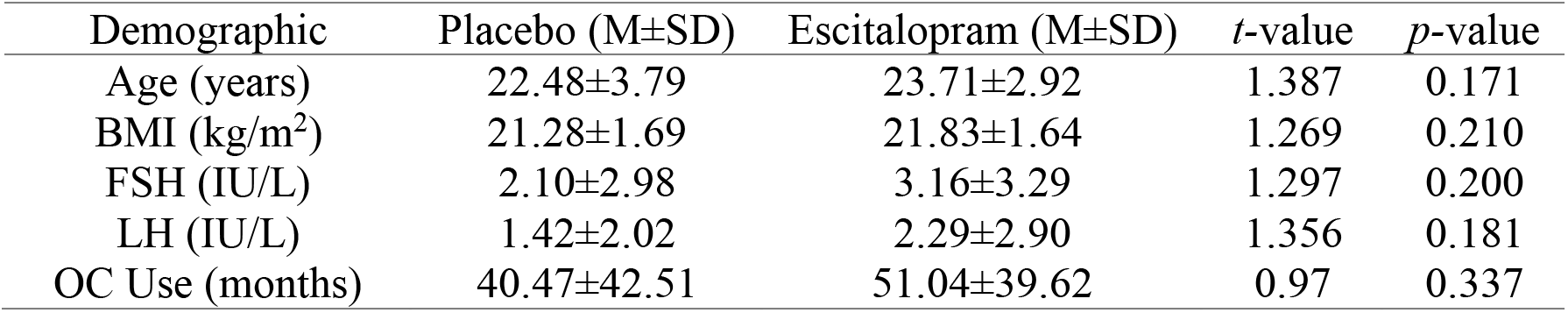
Group comparisons for baseline demographic characteristics. Results from independent sample *t*-tests assessing potential group differences for participant age, body mass index (BMI), Follicle stimulating hormone (FSH) levels, Luteinizing hormone (LH) levels, Length of oral contraceptive (OC) use. Mean±Standard Deviation (M±SD).

| Demographic | Placebo (M±SD) | Escitalopram (M±SD) | *t*-value | *p*-value |
| --- | --- | --- | --- | --- |
| Age (years) | 22.48±3.79 | 23.71±2.92 | 1.387 | 0.171 |
| BMI (kg/m<sup>2</sup>) | 21.28±1.69 | 21.83±1.64 | 1.269 | 0.210 |
| FSH (IU/L) | 2.10±2.98 | 3.16±3.29 | 1.297 | 0.200 |
| LH (IU/L) | 1.42±2.02 | 2.29±2.90 | 1.356 | 0.181 |
| OC Use (months) | 40.47±42.51 | 51.04±39.62 | 0.97 | 0.337 |

Analysis of STAI-S, POMS, and ESS questionnaires did not show group differences at single dose (STAI-S *t* = −1.76, *p* = 0.083; POMS *t* = 1.73, *p* = 0.088; ESS *t* =0.01, *p* = 0.987) or steady state (STAI-S *t* = −0.06, *p* = 0.945; POMS *t* = 1.07, *p* = 0.287; ESS *t* =0.23, *p* = 0.815). ASEC scores showed a significant group difference at single dose (*t* = −3.39, *p* = 0.002) but not steady state (*t* = −0.61, *p* = 0.551). However, we did not observe any associations between ASEC scores and plasma escitalopram levels (*R* = −0.29, *p* = 0.131), mean 1/f slope (*R* = 0.08, *p* = 0.660), or alpha power (*R* = 0.24, *p* = 0.228) at single dose in the escitalopram group. Moreover, we did not observe any associations between plasma levels and mean 1/f slope (*R* = −0.21, *p =* 0.282) or alpha power (*R* = 0.13, *p* = 0.503) at single dose.

### 3.2 Analysis of rsEEG

The intraclass correlation coefficient in the placebo group across the three timepoints was 0.923 for 1/f slope, 0.990 for power of alpha oscillations, and 0.998 for the conventional analysis of relative alpha power, suggesting excellent test-retest reliability. Analysis of 1/f slope yielded an effect of time, of group, and a group × time interaction (**Figure 2, Table 2**). We did not observe any significant interaction effect for power of alpha oscillations. Thus, post-hoc analyses and cluster-based permutations were only performed for 1/f slope.

**Figure 2.**
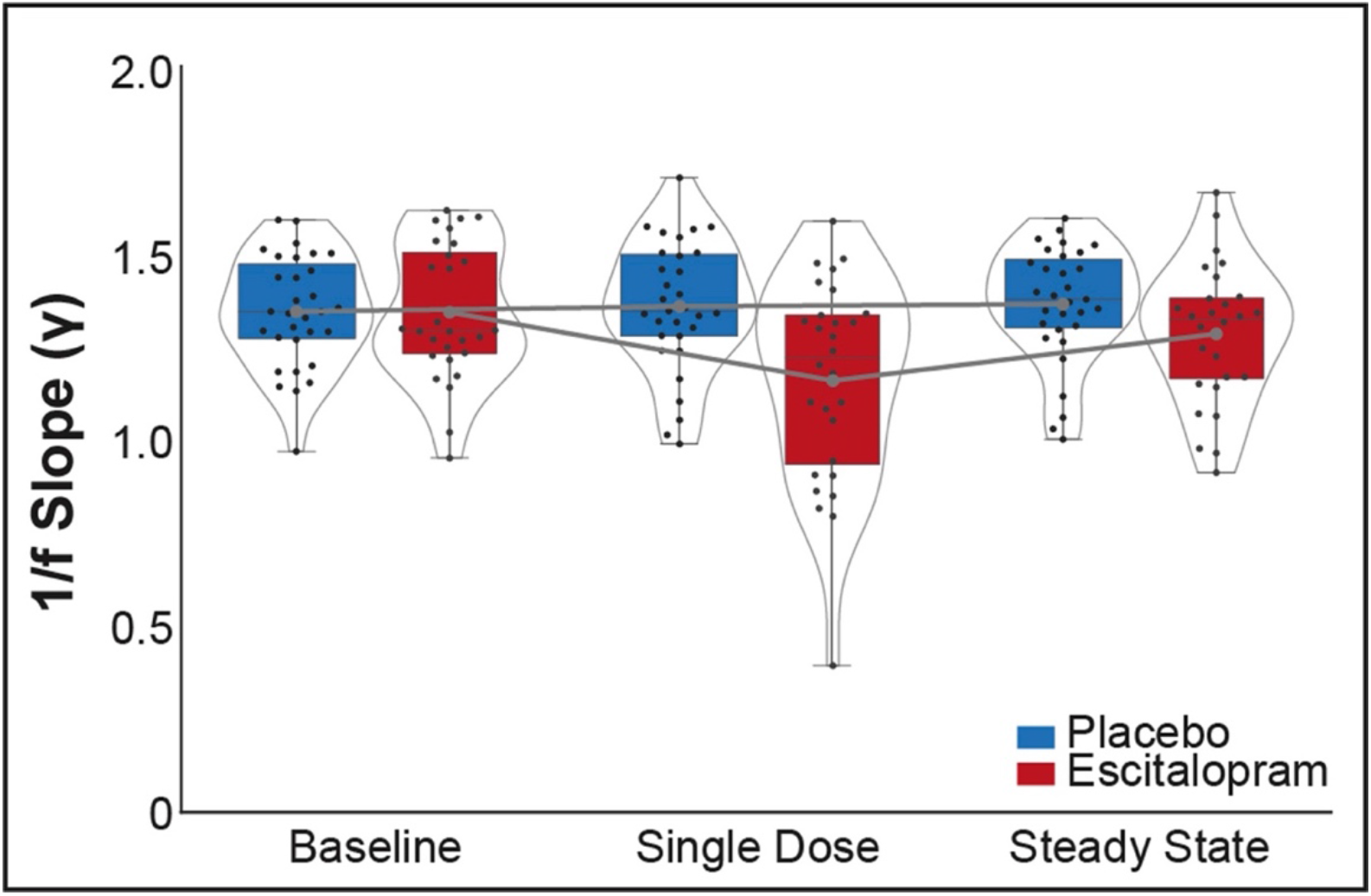
Significant changes in 1/f slope during one week of escitalopram administration. Linear mixed effects analysis shows that 1/f slope, the aperiodic component of the power spectral density, decreases following a single dose of escitalopram. Increases in 1/f slope are observed between single dose and steady state in the escitalopram group. Shown here are the individual data points (black dots) and mean values per group (gray dots). Inner box plot includes median and interquartile ranges, with whiskers extending 1.5 times the interquartile range. Width of the kernel densities reflects proportion of data located there.

**Table 2.** Linear mixed effects modeling of 1/f slope, detrended *α* power and relative *α* power. Results show omnibus effects of each fixed effect separately, with corresponding *p*-value and associated effect sizes. LRT = likelihood ratio test, df = degrees of freedom, χ^2^ = Chi-square, α = alpha. *significant contribution to model.

| Model Specification | Fixed Effects | LRT | | Marginal $R^2$ | Conditional $R^2$ |
| --- | --- | --- | --- | --- | --- |
| | | $\chi^2$ (df) | $p$-value | | |
| *1/f Slope* |  |  |  |  |  |
| Intercept | - | - | - | 0 | 0.374 |
| Time | time | 117.08 (2) | < 0.001* | 0.015 | 0.390 |
| Group | group+time | 4.51 (1) | 0.03* | 0.043 | 0.390 |
| Interaction | group*time | 169.40 (2) | < 0.001* | 0.064 | 0.411 |
| *$\alpha$ Power* |  |  |  |  |  |
| Intercept | - | - | - | 0 | 0.687 |
| Time | time | 1.18 (2) | 0.55 | 0.00 | 0.687 |
| Group | group+time | 3.31 (1) | 0.06 | 0.037 | 0.687 |
| Interaction | group*time | 4.53 (2) | 0.10 | 0.038 | 0.687 |
| *Relative $\alpha$ Power* |  |  |  |  |  |
| Intercept | - | - | - | 0 | 0.680 |
| Time | time | 30.90 (2) | <0.001* | 0.002 | 0.682 |
| Group | group+time | 4.04 (1) | 0.04 | 0.049 | 0.682 |
| Interaction | group*time | 55.03 (2) | <0.001* | 0.053 | 0.686 |

Post-hoc analyses for 1/f slope showed a significant group difference at single dose (*W* = −237, *p* = 0.002) but not steady state *W*=309, *p* = 0.058). Within groups comparisons over time show a significant time effect within the escitalopram group from (i) baseline to single dose (*V*=362, *p*<0.001), (ii) baseline to steady state (*V*=292, *p*=0.042), and (iii) from single dose to steady state (*V*=50, *p*<0.001) with decreased 1/f slope from baseline to single dose and an increase from single dose to steady state.

### 3.3 Cluster-based permutations in 1/f slope

Cluster-based group comparisons of 1/f slope did not show differences at baseline. Between-group comparisons at single dose (mean *z_electrode_* = −2.849, *p* = 0.002) and steady state (mean *z_electrode_* = −2.473, *p* = 0.026) show a significant decrease in the escitalopram group compared to placebo (**Figure 3A**). Within-escitalopram group comparisons showed a significant decrease in 1/f slope from baseline to single dose (mean *z_electrode_* = −3.136, *p* < 0.001) and from baseline to steady state (mean *z_electrode_* = −2.693, *p* = 0.004) (**Figure 3B**). Comparisons of single dose to steady state showed a significant increase in 1/f slope (mean *z_electrode_* = 2.772, *p* < 0.001). Mean power spectra were plotted for the cluster (electrodes F3, FC3, FT7, T7) significant in all contrasts (**Figure 3C**).

**Figure 3.**
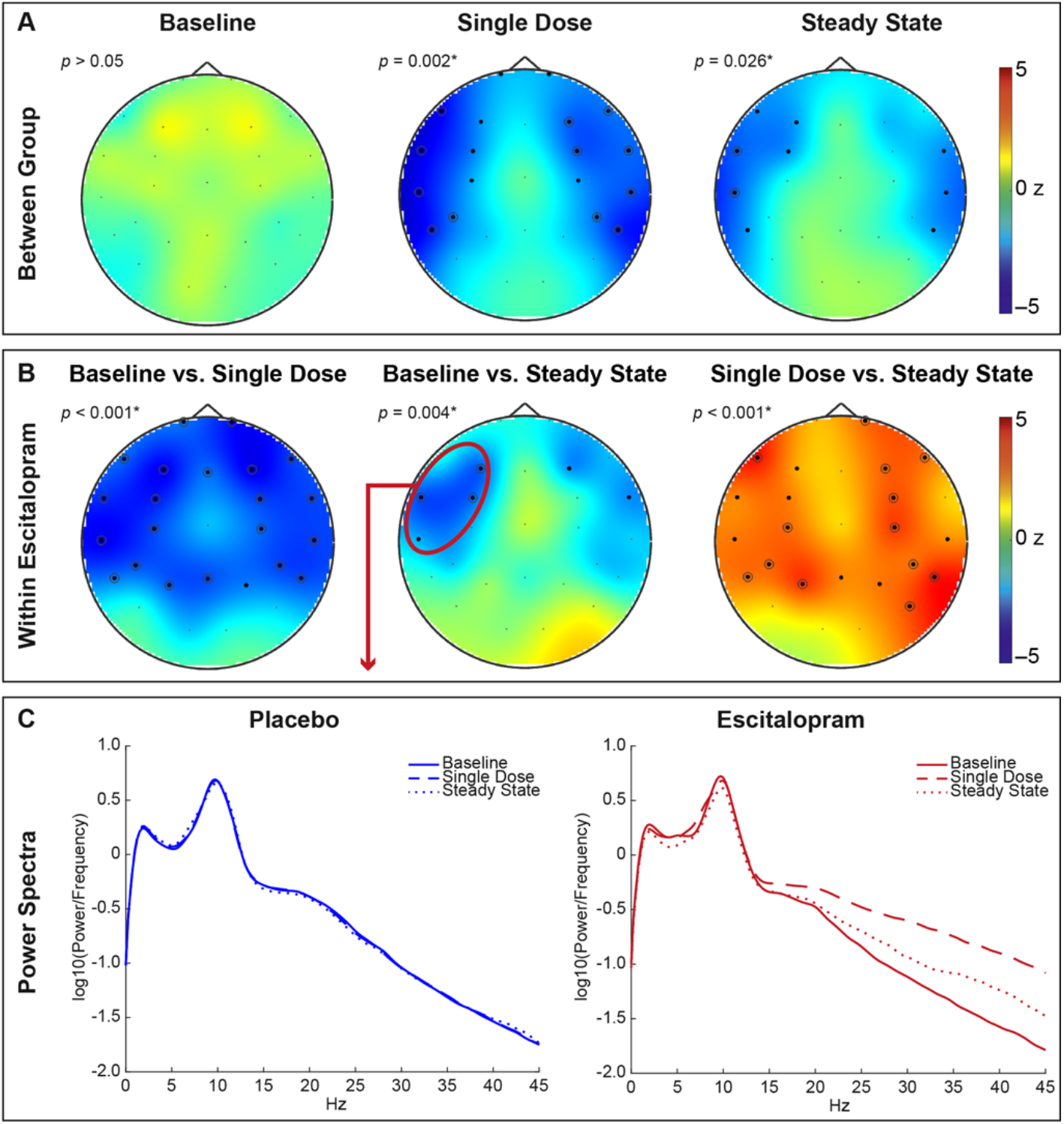
Cluster-based permutation tests show decreases in 1/f slope following escitalopram-intake at single dose and steady state. Shown are clusters surviving correction for multiple comparisons after computing 1,000 permutations. (**A**) We observe no group differences at baseline and significant clusters at single dose and steady state. (**B**) We observe significant clusters across all 3 assessments within the escitalopram group. *significant at *p* < 0.05, *p* = cluster statistic, *z* = effect size, ● = significant electrodes *p* < 0.05, ◉ = significant electrodes *p* < 0.01. (**C**) Mean power spectra plotted for cluster (electrodes F3, FC3, FT7, T7; indicated in red, Panel B) common to all significant clusters from permutation tests, in order to illustrate shifts in 1/f slope over one week of escitalopram-intake.

### 3.4 Linear regressions on residual 1/f slope in escitalopram group

Regression analyses on residual 1/f slope showed that baseline 1/f slope values did not predict single dose values (*R^2^_adj_* = −0.037, *p* = 0.849), baseline values predicted steady state values (*R^2^_adj_* = 0.235, *p* = 0.005), and single dose values predicted steady state values (*R^2^_adj_* = 0.462, *p* < 0.001) (**Figure 4A**).

**Figure 4.**
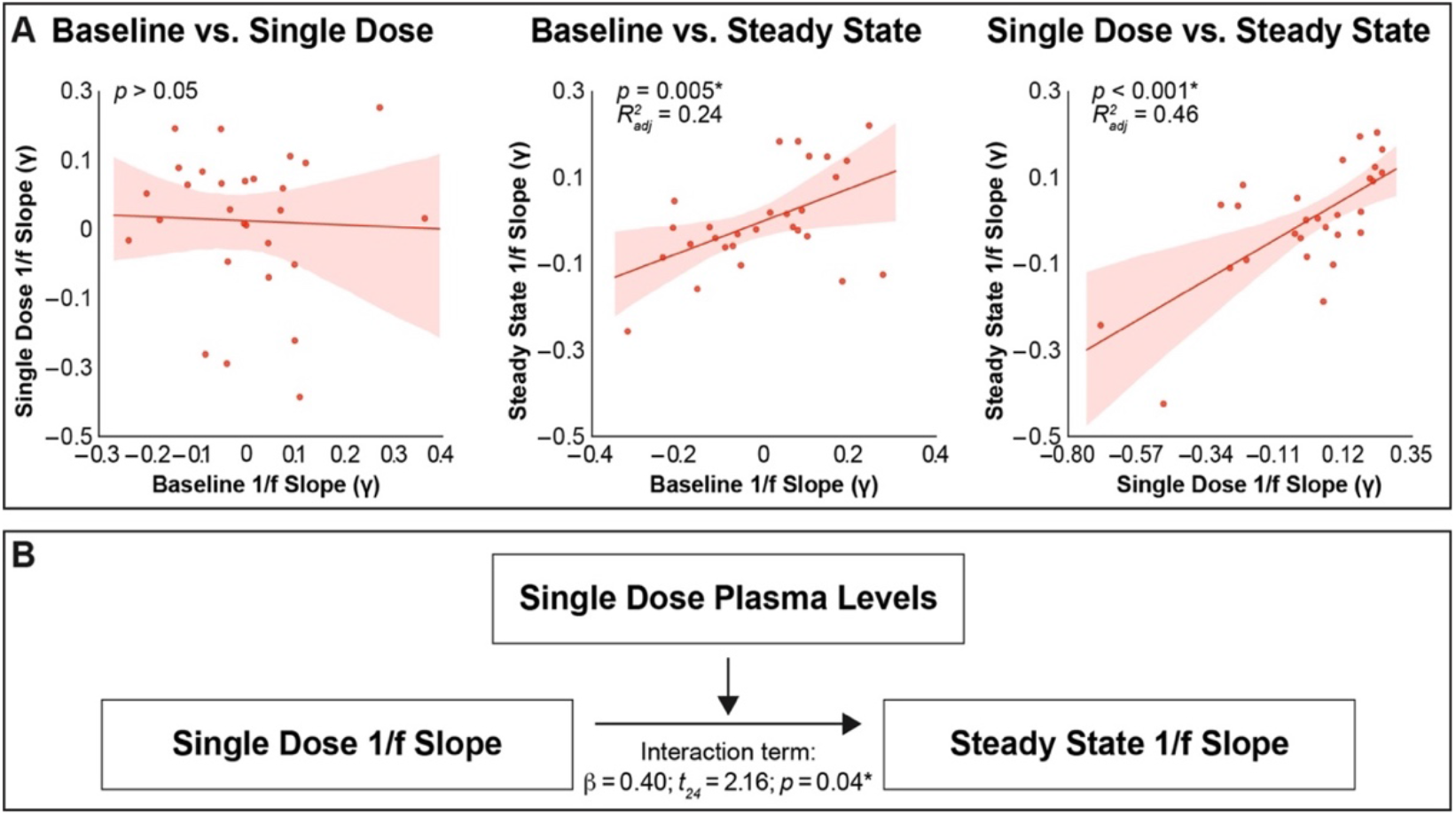
1/f slope at baseline and after a single dose of escitalopram predicts 1/f slope after one week of drug-intake. (**A**) Linear regression analyses of residuals (controlling for trait EEG signal) show that baseline 1/f slope predicts steady state response, and single dose 1/f slope predicts steady state response. Shaded area represents 95% confidence interval. (**B**) Moderation analysis shows that single dose plasma levels moderate the association between initial residual single dose response and maintained residual response at steady state. *significant at *p* < 0.05.

### 3.5 Moderation Analysis of residual 1/f slope in escitalopram group

For the exploratory moderation analysis, the overall model was significant (*F*_3,24_ = 11.16; *R*^2^ = 0.582; *p* < 0.001) (**Figure 4B**). The interaction of plasma and 1/f slope at single dose was significant (β = 0.40; *t*_24_ = 2.164; 95% BBCI 0.002 to 0.071; *p* = 0.04), suggesting that single dose plasma levels are a moderator of the association between the initial cortical response to escitalopram and the steady state response and associated with a maintained decrease in 1/f slope during SSRI intake.

### 3.6 Analysis of conventional relative alpha power

Relative alpha power analyses yielded an effect of time and a group × interaction (**Table 2**). Post-hoc analyses showed a group difference at single dose (*t* = 2.5, *p* = 0.013) but not one-week steady state (*t* = 1.7, *p* = 0.083).

## 4. Discussion

The present study reports changes in the aperiodic component of the PSD, a measure of cortical EIB, during steady state SSRI plasma levels in healthy female individuals. Our finding that one week of escitalopram-intake induces a widespread decrease of 1/f slope of the PSD, which represents an increase in EIB (32), suggests that escitalopram may tip the balance in favor of cortical excitability. Given that both baseline and single dose 1/f slope signals were associated with one-week steady state signal, and that escitalopram plasma levels after a single dose strengthened this relationship, we propose that 1/f slope could serve as a neurophysiological marker for individual cortical responsivity to SSRIs.

The significant decrease in 1/f slope following escitalopram administration is critical given the increasing focus on excitation-inhibition imbalance as a marker for neuropsychiatric disorders (1). While 1/f slope naturally changes with age in healthy humans (69), an exaggerated disruption of 1/f slope could represent noisier, less efficient neural circuits that contribute to plasticity-related deficits. Moreover, recent findings (70) demonstrated that the aperiodic component of the PSD is a more robust, stable measure of individual variability as compared to conventional power spectral features. Hence, our findings demonstrate that: (1) escitalopram reliably manipulates a stable measure of EIB, (2) 1/f slope at steady state levels of escitalopram can be predicted by baseline and single dose 1/f slope, and (3) inter-individual cortical responsivity can influence the strength of this relationship via escitalopram kinetics. Antidepressants have highly variable response rates in clinical settings (18), leading to weeks of trial and error (17). Our preclinical model identifies a neurophysiological indicator of individual SSRI cortical responsivity and thus establishes a framework to further characterize cortical responses to psychopharmacological intervention at a single-subject level in order to inform future translational research.

Due to the nature of rs-EEG, we cannot conclude if an increase in EIB following escitalopram administration is a result of increased excitation or decreased inhibition. Previous research suggests that the reversal of plasticity-related deficits depends on inhibitory transmission (2, 12, 13, 71). A recent review (13) proposes that SSRIs reactivate a plasticity period in the adult human brain by initially decreasing inhibitory tone, thus heightening cortical excitability. After multiples days of SSRI administration, a subsequent increase in inhibitory tone then re-establishes the balance, allowing for consolidation of these synaptic changes. We observe a similar pattern, with a widespread decrease in 1/f slope after a single dose of escitalopram, signifying an increasingly excitable state, followed by a slope increase from single dose to the one-week steady state assessment, with a spatially-confined region of decreased 1/f slope.

Unlike the previously mentioned studies that used SSRIs such as fluoxetine (12, 71–73), paroxetine (14), and sertraline (15), however, we administered the faster-acting SSRI escitalopram (38, 74, 75). Escitalopram has the highest degree of selectivity for binding to the serotonin transporter, thereby leading more quickly to increased extracellular serotonin levels (36, 37, 76). This increase activates various post-synaptic receptors that lead to complex interactions between the serotonergic, the inhibitory γ-aminobutyric acid (GABA)-ergic, and the excitatory glutamatergic systems (77). For example, escitalopram inhibits 5HT3 receptor currents *in vitro* (78), suggesting that escitalopram can enhance glutamate transmission by reducing GABA-mediated inhibition (77). Further support for escitalopram-induced increases in excitatory transmission comes from rodent models that have shown escitalopram enhances glutamate receptor subunit expression (79) and N-methyl-D-aspartate (NMDA)-mediated currents in rats (80), as well as hippocampal long-term potentiation (81). Thus, escitalopram may alter EIB through initially increasing excitatory transmission or decreasing inhibitory transmission. Future integration of quantitative neurochemical imaging, such as magnetic resonance spectroscopy to estimate the main excitatory and inhibitory neurotransmitters, would provide a more direct assessment that would be essential for understanding these dynamic changes in excitatory-inhibitory transmission across time, as we observe a relative increase followed by a decrease in EIB across the drug administration week. This subsequent decrease in EIB we find from single dose to one-week steady state is consistent with the view of functional serotonergic homeostasis underlying the adaptability of a healthy human adult brain (82, 83).

We also observe 1/f slope signal asymmetry at the one-week steady state assessment, with a spatially-confined region of significance in the left prefrontal hemisphere (**Figure 3B**). This finding is of interest in the context of previous studies (84, 85), which have also reported asymmetric findings in intrinsic brain activity following SSRI administration. Given the hypothesis that SSRIs stimulate neuroplasticity (10, 11, 86), a possible explanation for this observation is that frontal regions have been identified as a central hub in several cognitive (87), mood regulation (88), and memory processes (89, 90); and are thus essential to neuroplastic processes. For example, the left dorsolateral prefrontal cortex has been used as the target for transcranial magnetic stimulation (TMS) and neuromodulation in healthy controls (91–93), as well as the most efficacious and responsive target for the FDA-approved TMS-treatment for treatment resistant depression (94, 95). The specificity to the left hemisphere may also be attributable to an inherent asymmetry in the serotonergic system. While SSRIs block reuptake of serotonin by occupying the serotonin transporter, inhibitory 5HT1_A_ auto-receptor activation limits initial serotonin firing and release in cortical projection areas (96). Regional variation in this auto-inhibitory feedback mechanism, possibly due to an individual’s 5HT1_A_ auto-receptor density (82), distribution of 5HT1_A_ auto-receptors in the dorsal raphe nucleus versus whole brain hetero-receptors (97), or differences in hemispheric distribution of serotonin transporter or receptor density (98, 99), may serve as a trait-like signal that influences serotonin release and individual cortical responsivity to prolonged escitalopram (100, 101). In addition, we cannot exclude functional changes in the serotonergic system in response to escitalopram, such as shifts in receptor affinity. Such shifts have been shown to occur at the transcriptional level in a rodent model following sustained fluoxetine-administration (102) and could also contribute to the regional specificity of SSRI-effects. We acknowledge, however, the observational nature of this finding and future studies with direct quantification of inter-regional differences in the serotonergic system are required to discuss this interpretation in more detail.

Finally, we also report a significant relationship between neural responses to single dose administration and the neural responses to one-week escitalopram administration (**Figure 4A**), suggesting that initial neural responses to escitalopram may be informative for a steady state response in health. Given, however, that peripheral plasma escitalopram levels may be, to some degree, dissociated from brain kinetics (37, 103), we conducted our analysis viewing single dose plasma escitalopram levels as an early indicator of a peripheral bodily response to an SSRI. We therefore investigated whether these early peripheral pharmacokinetics (as reflected by plasma escitalopram levels following the first dose) interact with concurrent early neural responses to the drug (cortical EIB signal 1/f slope following the first dose) to predict the neural response to escitalopram after one week of intake (cortical EIB signal 1/f slope during relatively stable plasma levels). Our results show that single dose 1/f slope values and single dose plasma levels moderate the one-week steady state 1/f slope response. This finding suggests that early peripheral kinetics and the associated neural kinetics jointly influence the steady state neural response to escitalopram. While this finding advocates for the utility of 1/f slope as a metric of early pharmacologic sensitivity in both brain and body, we acknowledge that this explanation remains speculative and requires testing in specifically designed studies with larger and more diverse samples and in clinical populations, such as patients with depression or anxiety disorders.

Against our *a priori* hypotheses, we did not find group differences in power of alpha oscillations following SSRI intake. Unlike previous studies (23–26), however, we assessed alpha activity from the detrended PSD to avoid potential confounding effects of the broadband 1/f component. When we assessed relative alpha power without controlling for this component, we observed the hypothesized decrease in alpha power. These findings emphasize the importance of separately inspecting periodic oscillatory activity and aperiodic 1/f activity when testing narrowband oscillations such as alpha power, which have been shown to play functionally distinct roles (28). Our findings go beyond these previous studies, however, by extending this approach to a preclinical human model of SSRI-induced alterations in EIB.

One limitation of the study is that we investigated resting-state cortical EIB changes in a healthy population. We cannot infer how these changes would manifest in a clinical population, or how they would impact potential clinical outcome. However, changes in cortical excitability, similar to our observed effect, have been linked to changes in mood, attention, and cognitive performance in both healthy (28, 104) and patient populations (105, 106). Thus, while our findings do provide a model of escitalopram-induced changes in cortical EIB, future studies or existing datasets (107–109) should investigate whether SSRI-induced EIB changes early in treatment could predict outcomes in clinical environments. Secondly, replication studies are required to determine the generalizability of these findings to male participants, mid- and late-life populations, and naturally cycling female participants. Our sample of age-matched female participants using oral contraceptives was explicitly defined, however, to avoid potential confounding effects of sex and ovarian hormonal fluctuations on escitalopram responsivity (40, 46), resting-state connectivity (45, 47–49), and resting-state alpha activity specifically (50), as well as of age on 1/f slope (69). This is also an important demographic (45), as women of reproductive age often use oral contraceptives (110, 111), and oral contraceptive use has been associated with concurrent use of antidepressants (112). Thirdly, we acknowledge that there are certain limitations to this model, such as the assumption of desynchronized cortical states, and that there are alternative methodologies for non-invasive investigation of EIB (113, 114). We also acknowledge that changes in 1/f slope could have been driven by other physiological factors, such as mutual excitation among pyramidal cells (115) or arousal (29). While we cannot directly investigate the former, we can cautiously address the latter, as we observed no group nor time differences in the daytime sleepiness scale. Finally, we cannot speculate on potential dose-dependent effects or effects of other SSRIs, given our fixed 20 mg dose of escitalopram. This dose was chosen, however, as 20 mg reliably blocks 80 percent of the serotonin transporter (35–39).

## 5. Conclusions

By combining a novel measure to assess cortical EIB with a rigorously controlled interventional study design in health, we present first evidence, to our knowledge, for dynamic changes in EIB following one-week of escitalopram-intake. Moreover, our findings demonstrate the potential for 1/f slope as a neurophysiological marker for predicting individual cortical responsivity to SSRIs. Interventional studies in health are an important component in the decision-making process of whether to proceed to more comprehensive clinical trials in heterogenous patient populations. Given the continuously rising number of prescribed antidepressants (116), which are also more often prescribed to women (42), alongside the current underrepresentation of female samples in neuroscience research (43, 44), establishing this model in healthy female participants provides a timely framework to test the effects of a frequently prescribed SSRI on human cortical excitability. These findings provide a crucial stepping stone towards considering sex and hormone state in personalized treatment for depression and other neural-plasticity associated disorders.

## Acknowledgments

Preparation of this manuscript was supported by The Branco Weiss Fellowship – Society in Science, National Association for Research on Schizophrenia and Depression (NARSAD) Young Investigator Grant 25032 from the Brain & Behavior Research Foundation (awarded to Sacher), a Minerva Research Group grant from the Max Planck Society (awarded to Sacher), a Doctoral Scholarship from the FAZIT Foundation (awarded to Molloy), and a Fellowship from the Joachim Herz Foundation (awarded to Zsido).

## Data Availability

Data are available from the corresponding authors upon reasonable request R code publicly available at: https://github.com/EGGLab-2021/ZsidoMolloy2021_SSRI-EIB_R MatLab code publicly available at: https://github.com/EGGLab-2021/ZsidoMolloy2021_SSRI-EIB_Matlab

## References

1. Sohal VS, Rubenstein JL (2019): Excitation-inhibition balance as a framework for investigating mechanisms in neuropsychiatric disorders. Mol Psychiatry. 24:1248–1257.

2. Froemke RC (2015): Plasticity of cortical excitatory-inhibitory balance. Annu Rev Neurosci. 38:195–219.

3. Selten M, van Bokhoven H, Kasri NN (2018): Inhibitory control of the excitatory/inhibitory balance in psychiatric disorders. F1000Research. 7.

4. Rubenstein J, Merzenich MM (2003): Model of autism: increased ratio of excitation/inhibition in key neural systems. Genes, Brain and Behavior. 2:255–267.

5. Gao R, Penzes P (2015): Common mechanisms of excitatory and inhibitory imbalance in schizophrenia and autism spectrum disorders. Curr Mol Med. 15:146–167.

6. Molina JL, Voytek B, Thomas ML, Joshi YB, Bhakta SG, Talledo JA, et al. (2020): Memantine effects on electroencephalographic measures of putative excitatory/inhibitory balance in schizophrenia. Biol Psychiatry Cogn Neurosci Neuroimaging. 5:562–568.

7. Peterson EJ, Rosen BQ, Campbell AM, Belger A, Voytek B (2017): 1/f neural noise is a better predictor of schizophrenia than neural oscillations. Biorxiv. 113449.

8. Luscher B, Shen Q, Sahir N (2011): The GABAergic deficit hypothesis of major depressive disorder. Mol Psychiatry. 16:383–406.

9. Price RB, Duman R (2019): Neuroplasticity in cognitive and psychological mechanisms of depression: an integrative model. Mol Psychiatry. 1–14.

10. Castrén E (2005): Is mood chemistry? Nat Rev Neurosci. 6:241–246.

11. Castrén E (2013): Neuronal network plasticity and recovery from depression. JAMA psychiatry. 70:983–989.

12. Vetencourt JFM, Sale A, Viegi A, Baroncelli L, De Pasquale R, O’Leary OF, et al. (2008): The antidepressant fluoxetine restores plasticity in the adult visual cortex. Science. 320:385–388.

13. Schneider CL, Majewska AK, Busza A, Williams ZR, Mahon BZ, Sahin B (2019): Selective serotonin reuptake inhibitors for functional recovery after stroke: Similarities with the critical period and the role of experience-dependent plasticity. J Neurol. 1–7.

14. Gerdelat-Mas A, Loubinoux I, Tombari D, Rascol O, Chollet F, Simonetta-Moreau M (2005): Chronic administration of selective serotonin reuptake inhibitor (SSRI) paroxetine modulates human motor cortex excitability in healthy subjects. Neuroimage. 27:314–322.

15. Ilic TV, Korchounov A, Ziemann U (2002): Complex modulation of human motor cortex excitability by the specific serotonin re-uptake inhibitor sertraline. Neurosci Lett. 319:116–120.

16. Batsikadze G, Paulus W, Kuo M-F, Nitsche MA (2013): Effect of serotonin on paired associative stimulation-induced plasticity in the human motor cortex. Neuropsychopharmacology. 38:2260–2267.

17. Bschor T, Kern H, Henssler J, Baethge C (2018): Switching the antidepressant after nonresponse in adults with major depression: A systematic Literature search and meta-analysis.

18. Gaynes BN, Warden D, Trivedi MH, Wisniewski SR, Fava M, Rush AJ (2009): What did STAR* D teach us? Results from a large-scale, practical, clinical trial for patients with depression. Psychiatr Serv. 60:1439–1445.

19. Klimesch W, Sauseng P, Hanslmayr S (2007): EEG alpha oscillations: the inhibition–timing hypothesis. Brain research reviews. 53:63–88.

20. Pfurtscheller G (2001): Functional brain imaging based on ERD/ERS. Vision Res. 41:1257–1260.

21. Patat A, Trocherie S, Thébault J, Rosenzweig P, Dubruc C, Bianchetti G, et al. (1994): EEG profile of litoxetine after single and repeated administration in healthy volunteers. Br J Clin Pharmacol. 37:157.

22. Leuchter AF, Hunter AM, Jain FA, Tartter M, Crump C, Cook IA (2017): Escitalopram but not placebo modulates brain rhythmic oscillatory activity in the first week of treatment of major depressive disorder. J Psychiatr Res. 84:174–183.

23. Knott VJ, Howson AL, Perugini M, Ravindran AV, Young SN (1999): The effect of acute tryptophan depletion and fenfluramine on quantitative EEG and mood in healthy male subjects. Biol Psychiatry. 46:229–238.

24. Dumont G, De Visser S, Cohen A, Van Gerven J (2005): Biomarkers for the effects of selective serotonin reuptake inhibitors (SSRIs) in healthy subjects. Br J Clin Pharmacol. 59:495–510.

25. Olbrich S, Arns M (2013): EEG biomarkers in major depressive disorder: discriminative power and prediction of treatment response. International Review of Psychiatry. 25:604–618.

26. Kemp A, Griffiths K, Felmingham K, Shankman SA, Drinkenburg W, Arns M, et al. (2010): Disorder specificity despite comorbidity: resting EEG alpha asymmetry in major depressive disorder and post-traumatic stress disorder. Biol Psychol. 85:350–354.

27. Donoghue T, Haller M, Peterson EJ, Varma P, Sebastian P, Gao R, et al. (2020): Parameterizing neural power spectra into periodic and aperiodic components. Nat Neurosci. 23:1655–1665.

28. Ouyang G, Hildebrandt A, Schmitz F, Herrmann CS (2020): Decomposing alpha and 1/f brain activities reveals their differential associations with cognitive processing speed. Neuroimage. 205:116304.

29. Lendner JD, Helfrich RF, Mander BA, Romundstad L, Lin JJ, Walker MP, et al. (2020): An electrophysiological marker of arousal level in humans. Elife. 9:e55092.

30. Weber J, Klein T, Abeln V (2020): Shifts in broadband power and alpha peak frequency observed during long-term isolation. Scientific reports. 10:1–14.

31. Donoghue T, Dominguez J, Voytek B (2020): Electrophysiological frequency band ratio measures conflate periodic and aperiodic neural activity. Eneuro. 7.

32. Gao R, Peterson EJ, Voytek B (2017): Inferring synaptic excitation/inhibition balance from field potentials. Neuroimage. 158:70–78.

33. Colombo MA, Napolitani M, Boly M, Gosseries O, Casarotto S, Rosanova M, et al. (2019): The spectral exponent of the resting EEG indexes the presence of consciousness during unresponsiveness induced by propofol, xenon, and ketamine. Neuroimage. 189:631–644.

34. Robertson MM, Furlong S, Voytek B, Donoghue T, Boettiger CA, Sheridan MA (2019): EEG power spectral slope differs by ADHD status and stimulant medication exposure in early childhood. J Neurophysiol. 122:2427–2437.

35. Rao N (2007): The clinical pharmacokinetics of escitalopram. Clin Pharmacokinet. 46:281–290.

36. Kasper S, Sacher J, Klein N, Mossaheb N, Attarbaschi-Steiner T, Lanzenberger R, et al. (2009): Differences in the dynamics of serotonin reuptake transporter occupancy may explain superior clinical efficacy of escitalopram versus citalopram. Int Clin Psychopharmacol. 24:119–125.

37. Klein N, Sacher J, Geiss-Granadia T, Mossaheb N, Attarbaschi T, Lanzenberger R, et al. (2007): Higher serotonin transporter occupancy after multiple dose administration of escitalopram compared to citalopram: an [123 I] ADAM SPECT study. Psychopharmacology (Berl). 191:333–339.

38. Kasper S, Spadone C, Verpillat P, Angst J (2006): Onset of action of escitalopram compared with other antidepressants: results of a pooled analysis. Int Clin Psychopharmacol. 21:105–110.

39. Klein N, Sacher J, Geiss-Granadia T, Attarbaschi T, Mossaheb N, Lanzenberger R, et al. (2006): In vivo imaging of serotonin transporter occupancy by means of SPECT and [123 I] ADAM in healthy subjects administered different doses of escitalopram or citalopram. Psychopharmacology (Berl). 188:263–272.

40. LeGates TA, Kvarta MD, Thompson SM (2019): Sex differences in antidepressant efficacy. Neuropsychopharmacology. 44:140–154.

41. Nishizawa S, Benkelfat C, Young S, Leyton M, Mzengeza Sd, De Montigny C, et al. (1997): Differences between males and females in rates of serotonin synthesis in human brain. Proceedings of the National Academy of Sciences. 94:5308–5313.

42. Abbing-Karahagopian V, Huerta C, Souverein P, De Abajo F, Leufkens H, Slattery J, et al. (2014): Antidepressant prescribing in five European countries: application of common definitions to assess the prevalence, clinical observations, and methodological implications. Eur J Clin Pharmacol. 70:849–857.

43. Will TR, Proaño SB, Thomas AM, Kunz LM, Thompson KC, Ginnari LA, et al. (2017): Problems and progress regarding sex bias and omission in neuroscience research. eneuro. 4.

44. Beery AK, Zucker I (2011): Sex bias in neuroscience and biomedical research. Neuroscience & Biobehavioral Reviews. 35:565–572.

45. Taylor CM, Pritschet L, Jacobs EG (2020): The scientific body of knowledge–Whose body does it serve? A spotlight on oral contraceptives and women’s health factors in neuroimaging. Front Neuroendocrinol. 100874.

46. Barth C, Villringer A, Sacher J (2015): Sex hormones affect neurotransmitters and shape the adult female brain during hormonal transition periods. Frontiers in neuroscience. 9:37.

47. Pritschet L, Santander T, Taylor CM, Layher E, Yu S, Miller MB, et al. (2020): Functional reorganization of brain networks across the human menstrual cycle. Neuroimage. 220:117091.

48. Lisofsky N, Mårtensson J, Eckert A, Lindenberger U, Gallinat J, Kühn S (2015): Hippocampal volume and functional connectivity changes during the female menstrual cycle. Neuroimage. 118:154–162.

49. Petersen N, Kilpatrick LA, Goharzad A, Cahill L (2014): Oral contraceptive pill use and menstrual cycle phase are associated with altered resting state functional connectivity. Neuroimage. 90:24–32.

50. Brötzner CP, Klimesch W, Doppelmayr M, Zauner A, Kerschbaum HH (2014): Resting state alpha frequency is associated with menstrual cycle phase, estradiol and use of oral contraceptives. Brain Res. 1577:36–44.

51. Wittchen H-U, Wunderlich U, Gruschwitz S, Zaudig M (1997): SKID I. Strukturiertes Klinisches Interview für DSM-IV. Achse I: Psychische Störungen. Interviewheft und Beurteilungsheft. Eine deutschsprachige, erweiterte Bearb. d. amerikanischen Originalversion des SKID I.

52. Fydrich T, Renneberg B, Schmitz B, Wittchen H-U (1997): SKID II. Strukturiertes Klinisches Interview für DSM-IV, Achse II: Persönlichkeitsstörungen. Interviewheft. Eine deutschspeachige, erw. Bearb. d. amerikanischen Originalversion d. SKID-II von: MB First, RL Spitzer, M. Gibbon, JBW Williams, L. Benjamin,(Version 3/96).

53. Hampson E (2020): A brief guide to the menstrual cycle and oral contraceptive use for researchers in behavioral endocrinology. Horm Behav. 119:104655.

54. Molloy EN, Mueller K, Beinhoelzl N, Bloechl M, Piecha FA, Pampel A, et al. (2020): Modulation of premotor cortex response to sequence motor learning during escitalopram intake. J Cereb Blood Flow Metab.0271678X20965161.

55. Søgaard B, Mengel H, Rao N, Larsen F (2005): The pharmacokinetics of escitalopram after oral and intravenous administration of single and multiple doses to healthy subjects. The Journal of Clinical Pharmacology. 45:1400–1406.

56. Drewes P, Thijssen I, Mengel H (2001): A single-dose cross-over pharmacokinetic study comparing racemic citalopram (40 mg) with the s-enantiomer of citalopram (escitalopram, 20 mg) in healthy male volunteers. Poster presented at the Annual Meeting of the New Clinical Data Evaluation Unit (NCDEU), Phoenix (AZ) USA.

57. Spielberger CD (1983): Manual for the State-Trait Anxiety Inventory STAI (form Y)(“ self-evaluation questionnaire”).

58. Laux L, Glanzmann P, Schaffner P, Spielberger CD (1981): Das state-trait-angstinventar [The state-trait anxiety inventory]. Hogrefe, Göttingen (in German).

59. Dalbert C (2002): ASTS-Aktuelle Stimmungsskala.

60. Bloch KE, Schoch OD, Zhang JN, Russi EW (1999): German version of the Epworth sleepiness scale. Respiration. 66:440–447.

61. Uher R, Farmer A, Henigsberg N, Rietschel M, Mors O, Maier W, et al. (2009): Adverse reactions to antidepressants. Br J Psychiatry. 195:202–210.

62. Teichert J, Rowe JB, Ersche KD, Skandali N, Sacher J, Aigner A, et al. (2020): Determination of atomoxetine or escitalopram in human plasma by HPLC: Applications in neuroscience research studies. Int J Clin Pharmacol Ther.

63. Oostenveld R, Praamstra P (2001): The five percent electrode system for high-resolution EEG and ERP measurements. Clin Neurophysiol. 112:713–719.

64. Onton J, Makeig S (2006): Information-based modeling of event-related brain dynamics. Prog Brain Res. 159:99–120.

65. Bell AJ, Sejnowski TJ (1995): An information-maximization approach to blind separation and blind deconvolution. Neural Comput. 7:1129–1159.

66. RCore T (2013): R: A Language and Environment for Statistical Computing. R Foundation for Statistical Computing [Internet]. Vienna, Austria.

67. Maris E, Oostenveld R (2007): Nonparametric statistical testing of EEG-and MEG-data. J Neurosci Methods. 164:177–190.

68. Hayes AF (2017): Introduction to mediation, moderation, and conditional process analysis: A regression-based approach. 2nd ed.: Guilford Publications.

69. Voytek B, Kramer MA, Case J, Lepage KQ, Tempesta ZR, Knight RT, et al. (2015): Age-related changes in 1/f neural electrophysiological noise. J Neurosci. 35:13257–13265.

70. Demuru M, Fraschini M (2020): EEG fingerprinting: Subject-specific signature based on the aperiodic component of power spectrum. Comput Biol Med. 103748.

71. Pinto CB, Saleh Velez FG, Lopes F, de Toledo Piza PV, Dipietro L, Wang QM, et al. (2017): SSRI and motor recovery in stroke: reestablishment of inhibitory neural network tonus. Frontiers in neuroscience. 11:637.

72. Licinio A, Wong M-L, Licinio J (2017): Biological and behavioural antidepressant treatment responses with the selective serotonin reuptake inhibitor fluoxetine can be determined by the environment. Mol Psychiatry. 22:484–484.

73. Alboni S, Van Dijk RM, Poggini S, Milior G, Perrotta M, Drenth T, et al. (2017): Fluoxetine effects on molecular, cellular and behavioral endophenotypes of depression are driven by the living environment. Mol Psychiatry. 22:552–561.

74. Montgomery SA, Baldwin DS, Blier P, Fineberg NA, Kasper S, Lader M, et al. (2007): Which antidepressants have demonstrated superior efficacy? A review of the evidence. Int Clin Psychopharmacol. 22:323–329.

75. Montgomery SA, Möller H-J (2009): Is the significant superiority of escitalopram compared with other antidepressants clinically relevant? Int Clin Psychopharmacol. 24:111–118.

76. Sanchez C, Reines EH, Montgomery SA (2014): A comparative review of escitalopram, paroxetine, and sertraline: are they all alike? Int Clin Psychopharmacol. 29:185.

77. Pehrson AL, Sanchez C (2014): Serotonergic modulation of glutamate neurotransmission as a strategy for treating depression and cognitive dysfunction. CNS spectrums. 19:121–133.

78. Park YS, Sung K-W (2019): Selective serotonin reuptake inhibitor escitalopram inhibits 5-HT3 receptor currents in NCB-20 cells. Korean J Physiol Pharmacol. 23:509.

79. Ryan B, Musazzi L, Mallei A, Tardito D, Gruber SH, El Khoury A, et al. (2009): Remodelling by early-life stress of NMDA receptor-dependent synaptic plasticity in a gene–environment rat model of depression. Int J Neuropsychopharmacol. 12:553–559.

80. Schilström B, Konradsson-Geuken Å, Ivanov V, Gertow J, Feltmann K, Marcus MM, et al. (2011): Effects of S-citalopram, citalopram, and R-citalopram on the firing patterns of dopamine neurons in the ventral tegmental area, N-methyl-D-aspartate receptor-mediated transmission in the medial prefrontal cortex and cognitive function in the rat. Synapse. 65:357–367.

81. Bhagya V, Srikumar B, Raju T, Rao BS (2011): Chronic escitalopram treatment restores spatial learning, monoamine levels, and hippocampal long-term potentiation in an animal model of depression. Psychopharmacology (Berl). 214:477–494.

82. Carhart-Harris R, Nutt D (2017): Serotonin and brain function: a tale of two receptors. J Psychopharmacol (Oxf). 31:1091–1120.

83. Blier P, de MONTIGNY C, Chaput Y (1987): Modifications of the serotonin system by antidepressant treatments: implications for the therapeutic response in major depression. J Clin Psychopharmacol. 7:24S–35S.

84. Arnone D, Wise T, Walker C, Cowen PJ, Howes O, Selvaraj S (2018): The effects of serotonin modulation on medial prefrontal connectivity strength and stability: a pharmacological fMRI study with citalopram. Prog Neuropsychopharmacol Biol Psychiatry. 84:152–159.

85. Schaefer A, Burmann I, Regenthal R, Arélin K, Barth C, Pampel A, et al. (2014): Serotonergic modulation of intrinsic functional connectivity. Curr Biol. 24:2314–2318.

86. Umemori J, Winkel F, Didio G, Llach Pou M, Castrén E (2018): iPlasticity: Induced juvenile-like plasticity in the adult brain as a mechanism of antidepressants. Psychiatry Clin Neurosci. 72:633–653.

87. Miller EK, Cohen JD (2001): An integrative theory of prefrontal cortex function. Annu Rev Neurosci. 24:167–202.

88. Ray RD, Zald DH (2012): Anatomical insights into the interaction of emotion and cognition in the prefrontal cortex. Neuroscience & Biobehavioral Reviews. 36:479–501.

89. Kumar S, Zomorrodi R, Ghazala Z, Goodman MS, Blumberger DM, Cheam A, et al. (2017): Extent of dorsolateral prefrontal cortex plasticity and its association with working memory in patients with Alzheimer disease. JAMA psychiatry. 74:1266–1274.

90. Engel AK, Fries P, Singer W (2001): Dynamic predictions: oscillations and synchrony in top–down processing. Nat Rev Neurosci. 2:704–716.

91. Martin DM, Liu R, Alonzo A, Green M, Player MJ, Sachdev P, et al. (2013): Can transcranial direct current stimulation enhance outcomes from cognitive training? A randomized controlled trial in healthy participants. Int J Neuropsychopharmacol. 16:1927–1936.

92. Nikolin S, Loo CK, Bai S, Dokos S, Martin DM (2015): Focalised stimulation using high definition transcranial direct current stimulation (HD-tDCS) to investigate declarative verbal learning and memory functioning. Neuroimage. 117:11–19.

93. Mulquiney PG, Hoy KE, Daskalakis ZJ, Fitzgerald PB (2011): Improving working memory: exploring the effect of transcranial random noise stimulation and transcranial direct current stimulation on the dorsolateral prefrontal cortex. Clin Neurophysiol. 122:2384–2389.

94. O’Reardon JP, Solvason HB, Janicak PG, Sampson S, Isenberg KE, Nahas Z, et al. (2007): Efficacy and safety of transcranial magnetic stimulation in the acute treatment of major depression: a multisite randomized controlled trial. Biol Psychiatry. 62:1208–1216.

95. Cantone M, Bramanti A, Lanza G, Pennisi M, Bramanti P, Pennisi G, et al. (2017): Cortical plasticity in depression: a neurochemical perspective from transcranial magnetic stimulation. ASN neuro. 9:1759091417711512.

96. Artigas F, Romero L, de Montigny C, Blier P (1996): Acceleration of the effect of selected antidepressant drugs in major depression by 5-HT1A antagonists. Trends Neurosci. 19:378–383.

97. Hahn A, Lanzenberger R, Wadsak W, Spindelegger C, Moser U, Mien L-K, et al. (2010): Escitalopram enhances the association of serotonin-1A autoreceptors to heteroreceptors in anxiety disorders. J Neurosci. 30:14482–14489.

98. Fink M, Wadsak W, Savli M, Stein P, Moser U, Hahn A, et al. (2009): Lateralization of the serotonin-1A receptor distribution in language areas revealed by PET. Neuroimage. 45:598–605.

99. Madalena E, Lopes SS, Almeida A, Sousa N, Leite-Almeida H (2020): Unmasking the relevance of hemispheric asymmetries–break on through (to the other side). Prog Neurobiol.101823.

100. Garcia-Garcia AL, Newman-Tancredi A, Leonardo ED (2014): P5-HT 1A receptors in mood and anxiety: recent insights into autoreceptor versus heteroreceptor function. Psychopharmacology (Berl). 231:623–636.

101. Richardson-Jones JW, Craige CP, Guiard BP, Stephen A, Metzger KL, Kung HF, et al. (2010): 5-HT1A autoreceptor levels determine vulnerability to stress and response to antidepressants. Neuron. 65:40–52.

102. Le Poul E, Boni C, Hanoun Nm, Laporte A-M, Laaris N, Chauveau J, et al. (2000): Differential adaptation of brain 5-HT1A and 5-HT1B receptors and 5-HT transporter in rats treated chronically with fluoxetine. Neuropharmacology. 39:110–122.

103. Meyer JH, Wilson AA, Sagrati S, Hussey D, Carella A, Potter WZ, et al. (2004): Serotonin transporter occupancy of five selective serotonin reuptake inhibitors at different doses: an [11C] DASB positron emission tomography study. Am J Psychiatry. 161:826–835.

104. Cardone P, Van Egroo M, Chylinski D, Narbutas J, Gaggioni G, Vandewalle G (2020): Increased cortical excitability but stable effective connectivity index during attentional lapses. Sleep.

105. Ostlund BD, Alperin BR, Drew T, Karalunas SL (2021): Behavioral and cognitive correlates of the aperiodic (1/f-like) exponent of the EEG power spectrum in adolescents with and without ADHD. Developmental cognitive neuroscience. 48:100931.

106. Ferrarelli F, Phillips ML (2021): Examining and Modulating Neural Circuits in Psychiatric Disorders With Transcranial Magnetic Stimulation and Electroencephalography: Present Practices and Future Developments. Am J Psychiatry.appi. ajp. 2020.20071050.

107. Ulke C, Tenke CE, Kayser J, Sander C, Böttger D, Wong LY, et al. (2019): Resting EEG measures of brain arousal in a multisite study of major depression. Clin EEG Neurosci. 50:3–12.

108. Ang Y-S, Bruder GE, Keilp JG, Rutherford A, Alschuler DM, Pechtel P, et al. (2020): Exploration of baseline and early changes in neurocognitive characteristics as predictors of treatment response to bupropion, sertraline, and placebo in the EMBARC clinical trial. Psychol Med. 1–9.

109. Trivedi MH, McGrath PJ, Fava M, Parsey RV, Kurian BT, Phillips ML, et al. (2016): Establishing moderators and biosignatures of antidepressant response in clinical care (EMBARC): rationale and design. J Psychiatr Res. 78:11–23.

110. Nations U (2015): Trends in Contraceptive Use Worldwide. Obtenido de The Department of Economic and Social Affairs.

111. Daniels K, Abma JC (2020): Current contraceptive status among women aged 15–49: United States, 2017–2019.

112. Skovlund CW, Mørch LS, Kessing LV, Lidegaard Ø (2016): Association of hormonal contraception with depression. JAMA psychiatry. 73:1154–1162.

113. Trakoshis S, Rocchi F, Canella C, You W, Chakrabarti B, Ruigrok AN, et al. (2020): Intrinsic excitation-inhibition imbalance affects medial prefrontal cortex differently in autistic men versus women. Elife. 9:e55684.

114. Bruining H, Hardstone R, Juarez-Martinez EL, Sprengers J, Avramiea A-E, Simpraga S, et al. (2020): Measurement of excitation-inhibition ratio in autism spectrum disorder using critical brain dynamics. Scientific Reports. 10:1–15.

115. Freeman WJ (2006): Origin, structure, and role of background EEG activity. Part 4: Neural frame simulation. Clin Neurophysiol. 117:572–589.

116. Iacobucci G (2019): NHS prescribed record number of antidepressants last year. Bmj. 364:l1508.

